# Evaluation of feeding behaviour traits to predict efficiency traits in pigs using partial least square regression

**DOI:** 10.1101/2020.11.13.381103

**Authors:** E. Ewaoluwagbemiga, G. Bee, C. Kasper

## Abstract

The improvement of efficiency traits, such as protein efficiency (PE), digestible energy efficiency (EnE) and lipid gain (LipG), are relevant given their associations with environmental pollution, cost of productions, and the quality of meat. However, these traits are difficult traits to measure and usually require slaughtering of pigs. Efficiency traits are complex, and several factors, such as genetic predisposition, feed composition, but also individual feeding behaviour may contribute to efficiency. The objective of this study was therefore to evaluate the potential of using feeding behaviour traits to predict efficiency traits under dietary protein restriction. A total of 587 Swiss Large White pigs, consisting of 312 females and 275 castrated males, had *ad libitum* access to feed and water, and were fed a protein-reduced diet (80% of recommended digestible protein and essential amino acids) from 22.5 ± 1.6 to 106.6 ± 4.6 kg BW. Individual feed intake was monitored and carcass composition (lean and fat mass) at slaughter was determined by dual-energy X-ray absorptiometry (DXA). The PE and EnE were calculated as the ratio of protein or energy in the carcass (estimated by DXA) to the total protein or energy consumed. Feeding behaviour traits monitored were daily feed intake (DFI; g/day), feed intake per meal (FIM; g/meal), number of daily meals (NDM; meals/day), duration of meal (DUM; min/meal), feeding rate (FR; g/min), and feeder occupation (FO; min/day). A partial least square (PLS) regression was used to predict PE, EnE and LipG from feeding behaviour traits, while including farrowing series (for PE only), age at slaughter and body weight at slaughter. Accuracy of PLS regression was assessed based on RMSE and R^2^ for calibration and validation sets, and on concordance correlation coefficient (CCC), which were estimated over 100 replicates of calibration and validation sets. Models with a number of latent variables of 5, 2 and 3 were identified as optimal for PE, EnE, and LipG, which explained 34.64%, 55.42% and 82.68% of the total variation in PE, EnE, and LipG, respectively. Significant CCC were found between predicted and observed values for PE (0.50), EnE (0.70), and LipG (0.90). In conclusion, individual feeding behaviour traits can better predict EnE and LipG than for PE under dietary protein restriction when fed *ad libitum*.

**Implications:** This study suggests that five feeding behaviour traits, which are automatically recorded via feeder stations in large numbers with little effort, together with body weight and age, may be used to predict protein efficiency, energy efficiency and lipid gain in Swiss Large White pigs receiving a protein reduced diet with considerable accuracy. This will allow for easy collection of large amounts of data on these traits for precision feeding and genetic selection strategies, especially when additional traits are added in the future to further improve accuracy.

## Introduction

There is an increasing demand for a more sustainable pig production, lower production costs and high-quality meat products. Oftentimes, in order to guarantee maximal growth performance, pigs are fed more protein than necessary, but less than 50% of the ingested dietary protein is converted into carcass muscle (Kasper et al., 2020a; Millet et al., 2018), thereby contributing to environmental pollution. In order to achieve a more sustainable pig production, the improvement of protein efficiency (PE) is crucial. In addition to the need to improve PE in pig production, other traits such as energy efficiency (EnE) and lipid gain (LipG) are also important considering the cost of production and the quality of meat. In contrast to classical efficiency traits, such as residual feed intake (RFI) and feed conversion ratio (FCR), PE, EnE and LipG provide more detailed information about single nutrients and digestible energy. Specific information about the efficiency with which pigs convert dietary protein, digestible energy and lipids into lean mass, carcass energy content and lipid gain will enable the development of precision feeding or genetic selection strategies to reduce the environmental footprint of meat production. However, specific efficiency traits are difficult to measure due to costs and time involved, and the necessity to anesthetize or slaughter the animals. For instance, PE, EnE and LipG need detailed information on the protein, digestible energy and lipid content of the feed, the amount of feed ingested, and the content of those compounds in the carcass. Since an individual animal’s behaviour may contribute to efficiency (Herd et al., 2004), we hypothesize that feeding behaviour traits may be used to predict efficiency of pigs. Feeding behaviours include, but are not limited to, the time spent eating per meal, feed intake per day, time spent eating per day, number of meals and feeding rate. The availability of automated feeding stations has enabled the recording of detailed feeding behaviour traits in pigs without effort (Maselyne et al., 2015), which is completely non-invasive and allows the evaluation and selection of animals during their lifetime without the need for slaughter. Thus, we aimed to investigate whether feeding behaviour traits, in combination with other routinely recorded information, such as age and live BW at slaughter and farrowing series could be used in predicting efficiency traits.

A number of studies have investigated the relationships between feeding behaviour traits and feed conversion ratio (FCR) and residual feed intake (RFI) in pigs and cattle. The studies suggest that efficient animals with low RFI spend less time feeding each day and fewer frequency of feeding per day (Durunna et al., 2011; Nkrumah et al., 2006). Kavlak and Uimari (2019) reported that daily feed intake has strong genetic correlations (0.73 – 0.89) with FCR and RFI, while other feeding behaviour traits had low to moderate genetic correlations with FCR and RFI. Carcò et al. (2018) reported partial correlations, corrected for the effects of feeding treatments, of feeding behaviour traits with protein gain (−0.04 to 0.41) and lipid gain (−0.07 to 0.43) for pigs within a BW range of 46 – 145 kg. Although these studies have reported that some relationships exist between feeding behaviour traits and feed efficiency, it is unclear how pigs at a lower BW range (20 – 100 kg) respond to a protein-reduced diet and how this feeding response may be used in predicting efficiency traits (PE, EnE, and LipG). The objective of this study was therefore to investigate the possibility of predicting efficiency traits (PE, EnE, and LipG) from automatically recorded feeding behaviour traits and other routinely recorded traits using partial least square (PLS) regression.

## Materials and Methods

### Animals

A total of 587 Swiss Large White pigs (N_females_ = 312, N_castrated-males_ = 275) from twelve farrowing series were used, which had *ad libitum* access to feed and water. Piglets were weaned on average at 27 ± 2 days after birth by removing the sow, after which they remained in the farrowing pen up to an average age of 28 ± 5 days and fed a starter diet, which was formulated according to the current Swiss feeding recommendations for swine and contained standard protein levels. Thereafter, pigs were placed in pens equipped with automatic feeders and stayed on the starter diet. Every week, pigs were weighed individually and, once the pig reached 20 kg (average BW of 22.5 ± 1.6 kg), it was allocated to a new pen. This was done until a maximum number of 12/24/48 pigs per pen (depending on the pen layout; minimum 1m^2^ per pig and maximum 12 pigs/feeder station) was reached. Pigs remained in this pen until slaughter. A total of 7 pens were used throughout the farrowing series; 4 pens were equipped with two single-spaced automatic feeder stations with individual pig recognition system (Schauer Maschinenfabrik GmbH & Co. KG, Prambachkirchen, Austria). Another 2 pens had 4 feeder stations each housed 24 pigs, and 1 pen had 8 feeder stations for 48 pigs. For each pen, half of the feeder stations dispensed grower diet until an average BW of 63 ± 2 kg, and the other half dispensed the finisher diet from an average BW of 63 ± 2 kg until an average BW of 106 ± 5 kg when the pigs were slaughtered. The automatic feeder stations recorded all daily visits to the feeder, time of feeding, duration of feeding and quantity of feed consumed per visit for each animal. The starter diet was formulated according to the current Swiss feeding recommendations for swine, whereas the levels of digestible protein and essential amino acids for the grower and finisher diets were 20% lower than the recommended levels (Agroscope, 2017). Hence, the grower and finisher diets contained on average 134.41 (± 3.55) g and 117.71 g CP (± 3.15) per kg feed, respectively, with the exact same digestible energy content (13.2 MJ/kg) as the recommended diet.

### Feeding behaviours and efficiency traits

The automated single-spaced feeder stations continuously recorded the date and time of each single feeder visit and the amount of feed eaten by each individual pig, from which a range of feeding behaviours were computed individually, and the average of each individual was used for further analysis. Visits separated by less than 5 min were grouped together and considered as meals (De Haer and Merks, 1992). Records of meals with no feed intake (i.e., 0 g) were removed (N = 9,617 meals). Records of meals with an unusual amount of feed ingested, i.e., less than 0.2 g/min, which average 1.5 g of feed in 12 min (N = 198 meals) and greater than 300 g/min, which average 1362 g of feed in 3 min (N = 4), were considered as physiologically impossible and were removed. After edits, 614,871 meals were recorded. Feeding behaviour traits evaluated were average daily feed intake (DFI; g/day), number of daily meals (NDM; meals/day), time spent feeding per day (FO; min/day), feed intake per meal (FIM = DFI/NDM; g/meal), duration per meal (DUM = FO/NDM; min/meal) and feeding rate (FR = DFI/FO; g/min).

Pigs were slaughtered at a BW of approximately 106 ± 5 kg and the left carcass including the whole head was scanned with dual-energy X-ray absorptiometry (DXA; GE Lunar i-DXA, GE Medical Systems, Glattbrugg, Switzerland), which determines, among other things, the lean tissue and fat content. The lean tissue and the fat content obtained from i-DXA was used in the following prediction equations to estimate the protein content, energy content and fat content retained in the carcass (Kasper et al., 2020b).

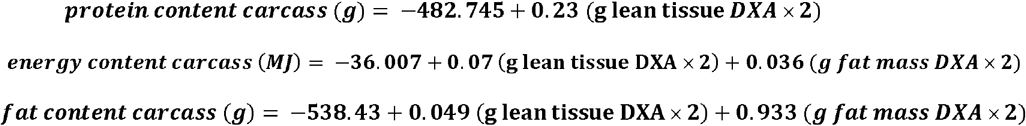

Protein and energy efficiency of the carcass was thereafter calculated as the ratio of protein or digestible energy retained in the carcass (corrected for protein or energy content in the carcass at 20 kg live BW) to the total protein or digestible energy intake.

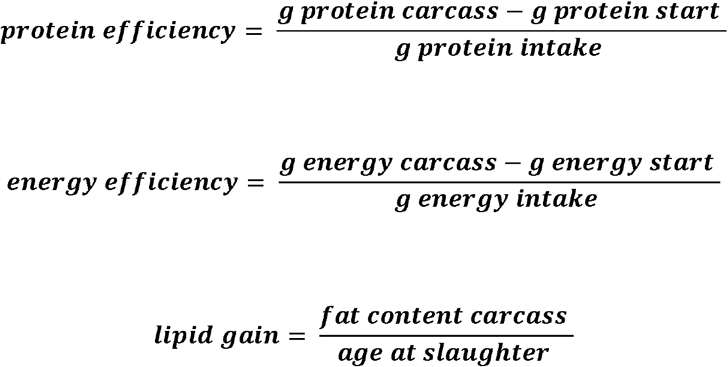

Protein content and energy content of pigs at the start of this experiment (***g protein start and g energy start***) was estimated from a sample of 24 piglets slaughtered at 21.26 ± 1.59 kg BW in a previous experiment (Ruiz-Ascacibar et al., 2017). The protein and digestible energy content per kg carcass of these 24 piglets were chemically determined and the average of each sex was used as the baseline protein and energy content of piglets at 20 kg BW. The protein content and energy content of pigs at the start of the present experiment were estimated by multiplying the actual live BW of pigs when the reduced-protein diet experiment started with the protein and energy content per kg carcass of piglet, respectively, as previously determined from the 24 piglets. The digestible energy intake of pig was calculated by multiplying the total amount of feed eaten from the start of the experiment until slaughter with the estimated digestible energy content of the diet (13.2 MJ/kg). The digestible energy of diet was estimated using the chemical composition of diet and digestibility coefficient in a regression equation. The digestibility coefficients were obtained from the Swiss Feed database (Agroscope, 2017).

### Statistical analysis

Data were analyzed with R software V 3.6.3. Two pigs with high protein and energy efficiency were considered as outliers from a boxplot and were therefore removed from the analysis. Both the feeding behaviour traits (except DFI) and efficiency traits (PE, EnE and LipG) showed deviation from normality as determined with the Shapiro-Wilk test. Feeding behaviour traits, except for DFI, were therefore log-transformed. However, based on visual inspection of the histogram of raw data and QQ-plots, the efficiency traits were considered as normally distributed. Due to differences in scales, feeding behaviour traits were scaled and centred to a mean of 0 and standard deviation of 1, likewise body weight and age at slaughter. However, efficiency traits were not scaled to allow easy interpretation of results.

Due to significant correlations between the feeding behaviour traits (Table 1), partial least square (PLS) regression was used for analysis, as this method can better handle collinearity between variables. The PLS regression was conducted using the package *pls* (Mevik et al., 2020) in a 4-step process. First, for each efficiency trait, an initial PLS model was run on the full dataset including all feeding behaviour traits as independent variables. Weight at slaughter, sex, age at slaughter, feeder station and farrowing series were also included to account for the effect of these variables on the efficiency traits. Thereafter, the data set was randomly divided into calibration and validation data sets, which contained 80% (469 pigs) and 20% (116 pigs) of the full data set, respectively. R^2^, RMSE, and concordance correlation coefficient (CCC) for each efficiency trait were estimated using 100 replicates of the calibration and validation sets. All variables were used as continuous variables in the PLS regression models, except for feeder station, sex and farrowing series, which were used as categorical variables. Only predictors with a variable of importance for projection (VIP), which is a score used for variable selection, greater than one, were used in the final models. Second, in the training step, a linear PLS regression model containing only the selected variables was fitted on the calibration data set using a leave-one-out (LOO) cross-validated prediction.

**Table 1.**
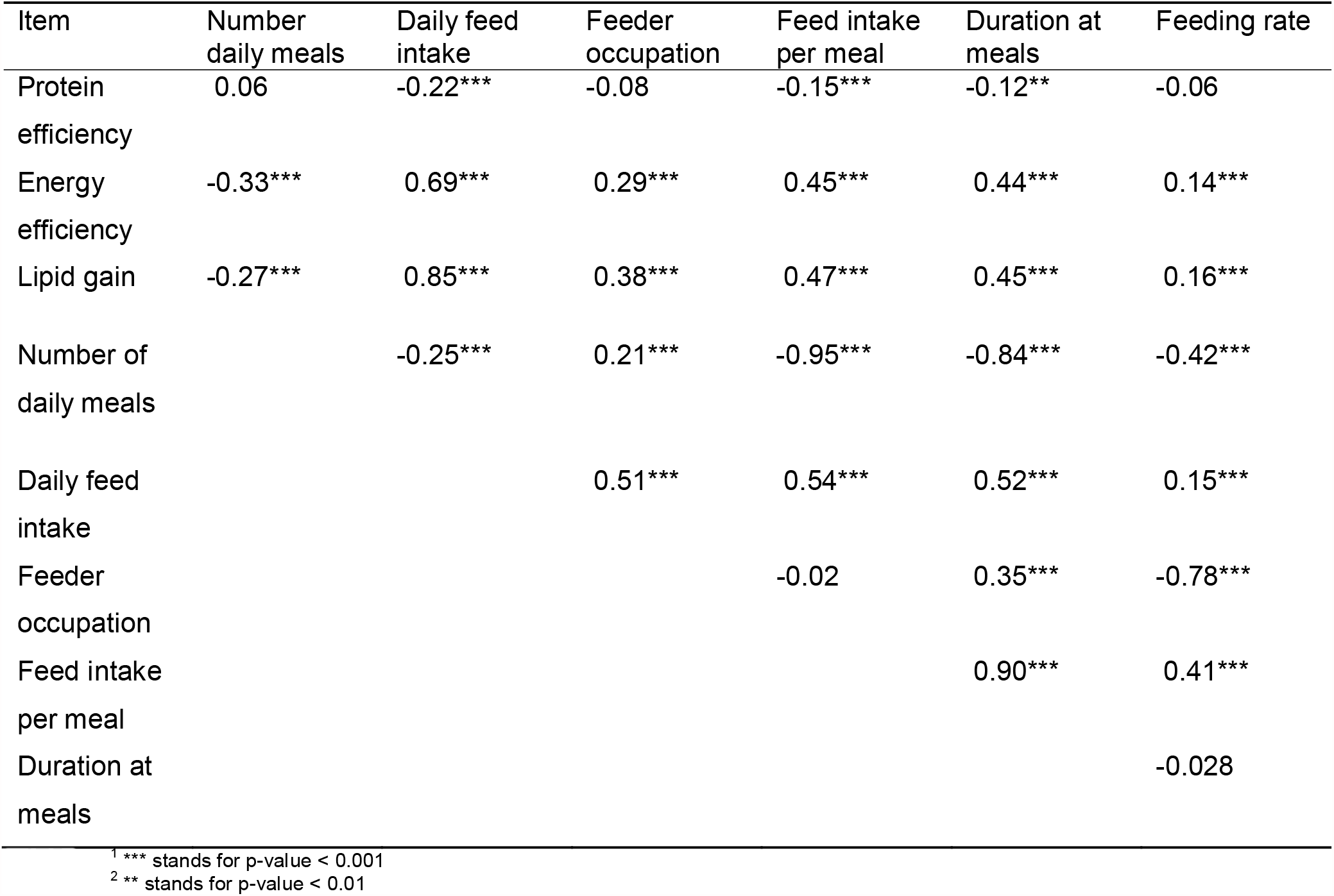
Pearson correlations among feeding behaviour traits^1^

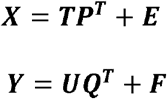

Where ***X*** is a design matrix for predictors (feeding behaviour traits, farrowing series, age at slaughter and body weight at slaughter with VIP > 1). ***Y*** is a matrix of response variables (protein efficiency, energy efficiency and fat retention); ***T*** and ***U*** are matrices that are projections of ***X*** and ***Y***, respectively; ***P*** and ***Q*** are orthogonal loading matrices, and ***E*** and ***F*** are the error terms. Third, the optimal number of components (latent variables; LV) retained was determined for each model by the so-called permutation approach implemented in the *pls* package, which tests whether adding new component to the model is beneficial (as described in the package vignette; Wherens and Mevik, 2007). Finally, in the validation step, the model obtained in the training step using the optimal number of LV determined previously was applied on the validation set. The accuracy of the PLS regression model was evaluated based on RMSE and coefficient of determination (R^2^) for calibration (R^2^c) and validation (R^2^v). The mean bias and a concordance correlation coefficient (CCC), which measures the agreement between two variables were used to assess the accuracy of prediction from the PLS models. To investigate the robustness of our results across statistical methods, we compared prediction accuracies between PLS regression and Bayesian models (Bayesian ridge regression, BayesA and BayesB; Supplementary Table S1). Results showed robustness and PLS regression was therefore used in the further analysis for this study.

## Results

The descriptive statistics for feeding behaviour traits and nutrient composition for Swiss Large White pigs are presented in Table 2. All feeding behaviour traits, except for FR, had VIP > 1 and were considered important in the models predicting the efficiency traits (Figure 1). Age and body weight at slaughter also had VIP > 1 for all the efficiency traits and were considered in the models (Figure 1). Since farrowing series 3 and 10 had VIP > 1 for PE, farrowing series was included in the model predicting PE. Thus, FIM, DFI, FO, NDM, DUM, Age and body weight at slaughter were included in the final models predicting the efficiency traits, and farrowing series in the model predicting PE. One standard deviation change in DFI, log-transformed FIM, log-transformed NDM, was associated with a 1.84%, 0.39%, 0.14% lower PE, while one standard deviation change in log-transformed DUM and FO was associated with 0.20% and 0.11% higher PE (Figure 2), respectively. One standard deviation change in DFI, log-transformed FIM, log-transformed FO and log-transformed DUM was associated with 0.84%, 0.33%, 0.27% and 0.21% higher EnE, respectively (Figure 3). One standard deviation change in DFI, log-transformed FIM, log-transformed FO and log-transformed DUM was associated with 8.01 g/day, 2.39 g/day, 0.47 g/day and 0.07 g/day higher LipG, respectively (Figure 4).

**Table 2.** Descriptive statistics of body weight, feeding behaviour traits and efficiency traits.

| Variables | Mean ± SD | Minimum | Maximum |
| --- | --- | --- | --- |
| **Body weight (kg)** |  |  |  |
| Start of grower period | 22.5 ± 1.6 | 18.9 | 27.3 |
| Start of finisher period | 63.5 ± 2.3 | 60 | 79.8 |
| Slaughter | 106.6 ± 4.6 | 88.6 | 120.8 |
| **Feeding behaviour traits** |  |  |  |
| DFI<sup>1</sup> (kg/day) | 2.31 ± 0.26 | 1.54 | 3.03 |
| NDM<sup>2</sup> (meals/day) | 10.19 ± 3.10 | 4.27 | 22.07 |
| FO<sup>3</sup> (min/day) | 61.02 ± 10.75 | 35.97 | 97.50 |
| FR<sup>4</sup> (g/min) | 38.69 ± 5.93 | 24.28 | 58.17 |
| DUM<sup>5</sup> (min/meal) | 6.50 ± 2.16 | 2.50 | 16.43 |
| FIM<sup>6</sup> (g/meal) | 251 ± 91 | 88.87 | 673.55 |
| **Efficiency traits** |  |  |  |
| Protein consumed (g) | 29088 ± 1805 | 24740 | 35078 |
| Digestible energy consumed (MJ) | 3309 ± 198 | 2784 | 3955 |
| Protein gain (g/day) | 78.46 ± 6.42 | 56.48 | 114.22 |
| Lipid gain (g/day) | 126 ± 20 | 64.01 | 197.23 |
| Energy gain (g/day) | 6.58 ± 0.86 | 3.76 | 9.50 |
| Protein retention (g) | 13702 ± 754 | 11242 | 20902 |
| Digestible energy retention (MJ) | 1236 ± 108 | 817 | 1665 |
| Protein efficiency | 0.38 ± 0.02 | 0.31 | 0.47 |
| Energy efficiency | 0.33 ± 0.03 | 0.23 | 0.42 |

<sup>1</sup> DFI: daily feed intake
<sup>2</sup> NDV: number of daily meals
<sup>3</sup> FO: feeder occupation
<sup>4</sup> FR: feeding rate
<sup>5</sup> DUV: duration of meal
<sup>6</sup> FIV: feed intake per meal

**Figure 1:**
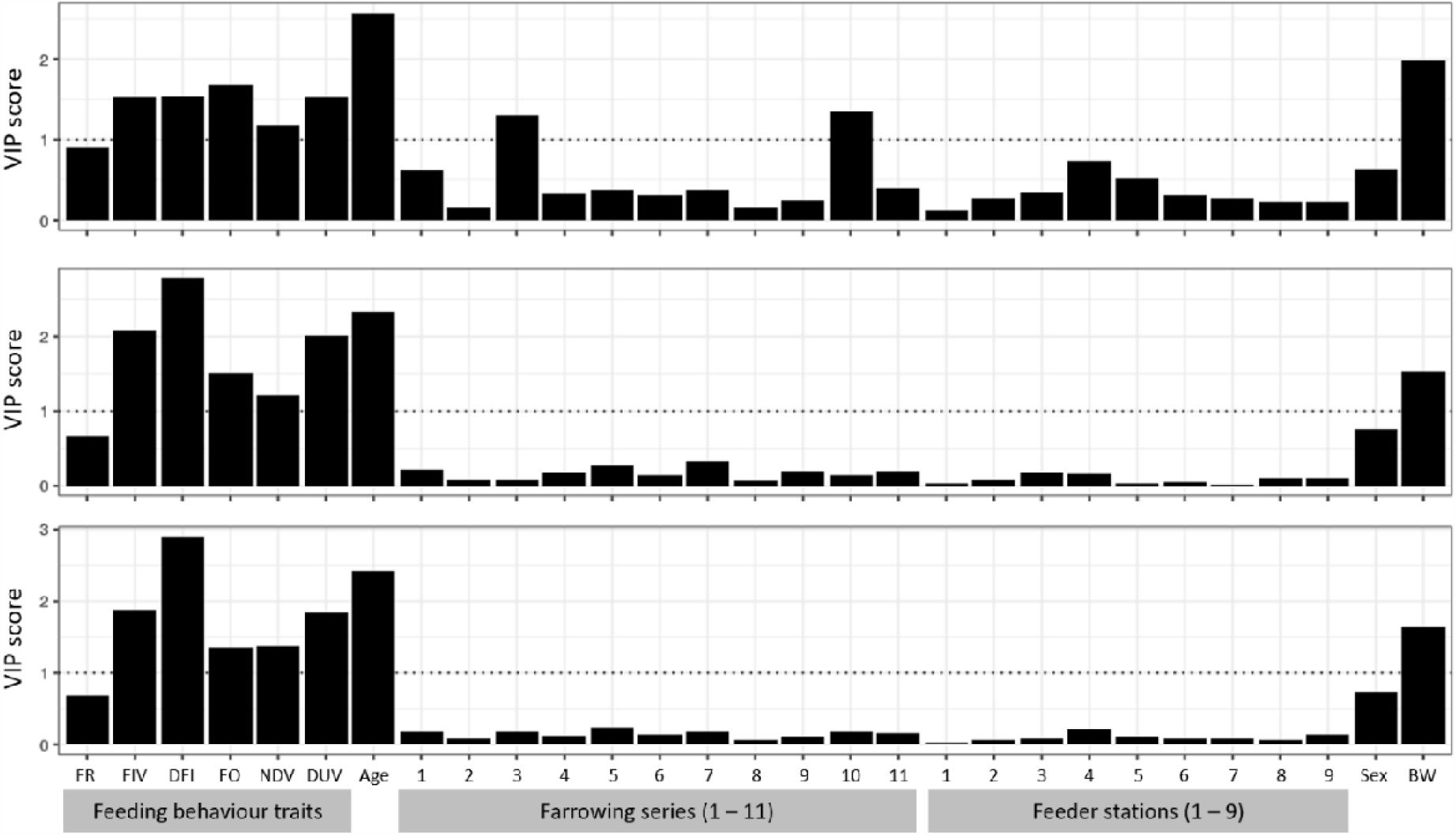
variable of importance for projection (VIP) for protein efficiency (top), energy efficiency (middle) and lipid gain (lower). FR: feeding rate, FIV: feed intake per visit, DFI: daily feed intake, FO: feeder occupation, NDV: number of daily visits, DUV: duration of visit, Age: age at slaughter, BW: body weight at slaughter

**Figure 2:**
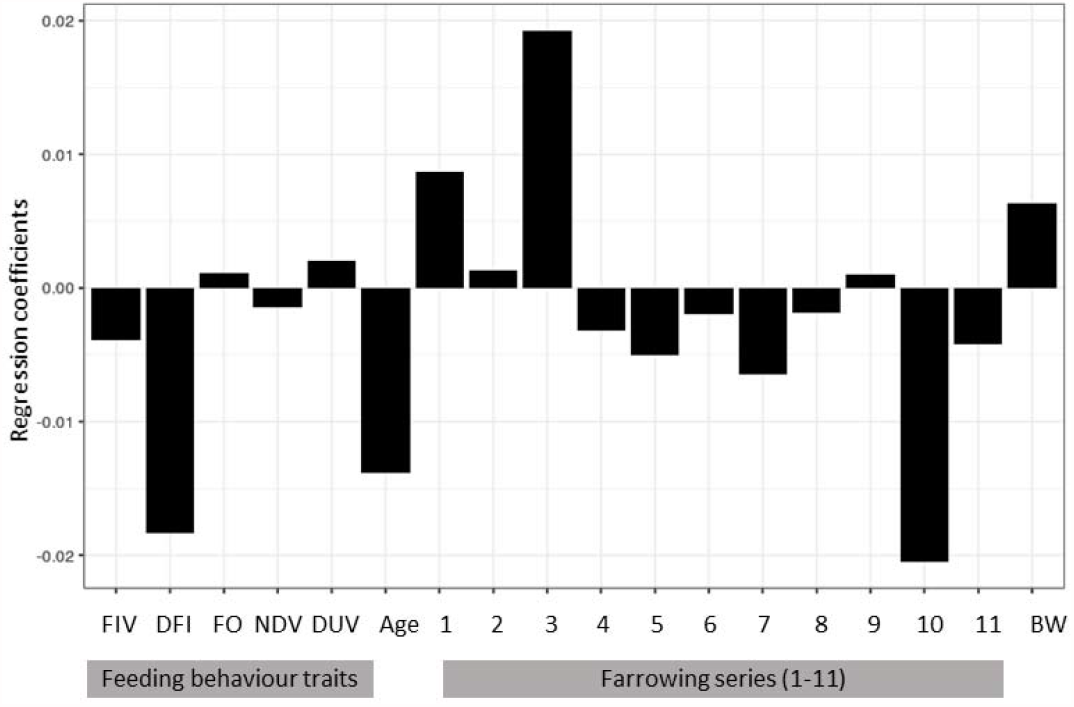
regression coefficient for daily feed intake, age at slaughter and body weight at slaughter and log-transformed regression coefficients for feed intake per visit, feeder occupation, number of daily visits and duration of visit from partial least square regression using feeding behaviour traits to predict protein efficiency, and including the effects of age at slaughter, farrowing series and body weight at slaughter. FIV: feed intake per visit, DFI: daily feed intake, FO: feeder occupation, NDV: number of daily visits, DUV: duration of visit, Age: age at slaughter, BW: body weight at slaughter

**Figure 3:**
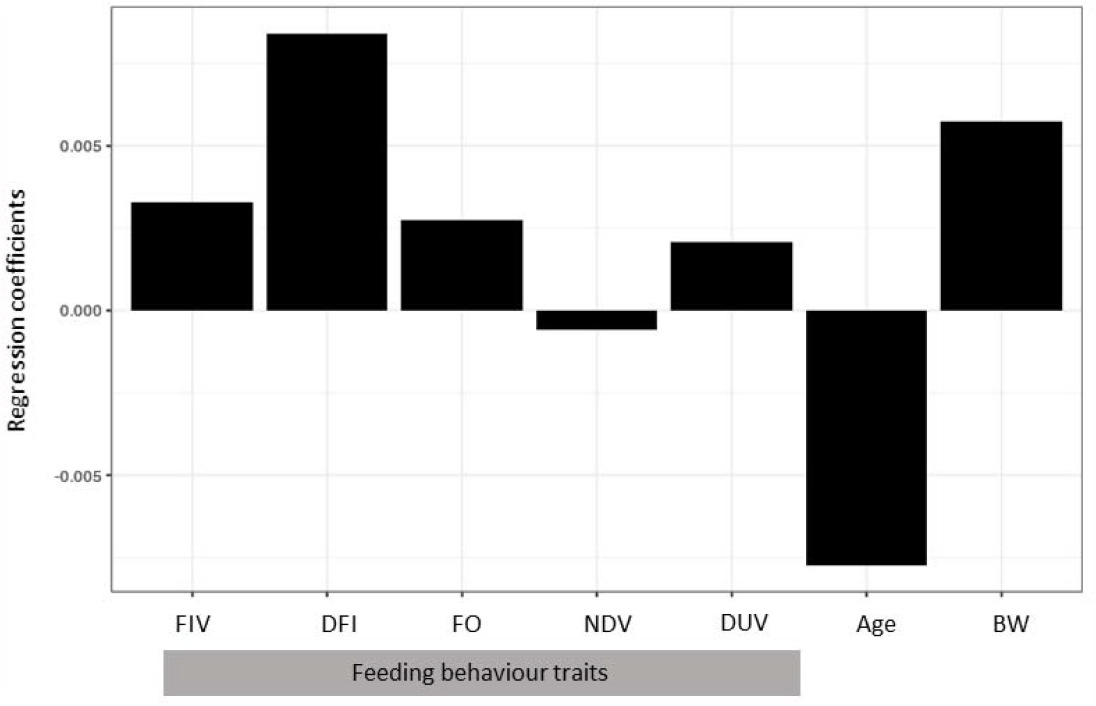
regression coefficient for daily feed intake, age at slaughter and body weight at slaughter and log-transformed regression coefficients for feed intake per visit, feeder occupation, number of daily visits and duration of visit from partial least square regression using feeding behaviour traits to predict energy efficiency, and including the effects of age at slaughter and body weight at slaughter. FIV: feed intake per visit, DFI: daily feed intake, FO: feeder occupation, NDV: number of daily visits, DUV: duration of visit, Age: age at slaughter, BW: body weight at slaughter

**Figure 4:**
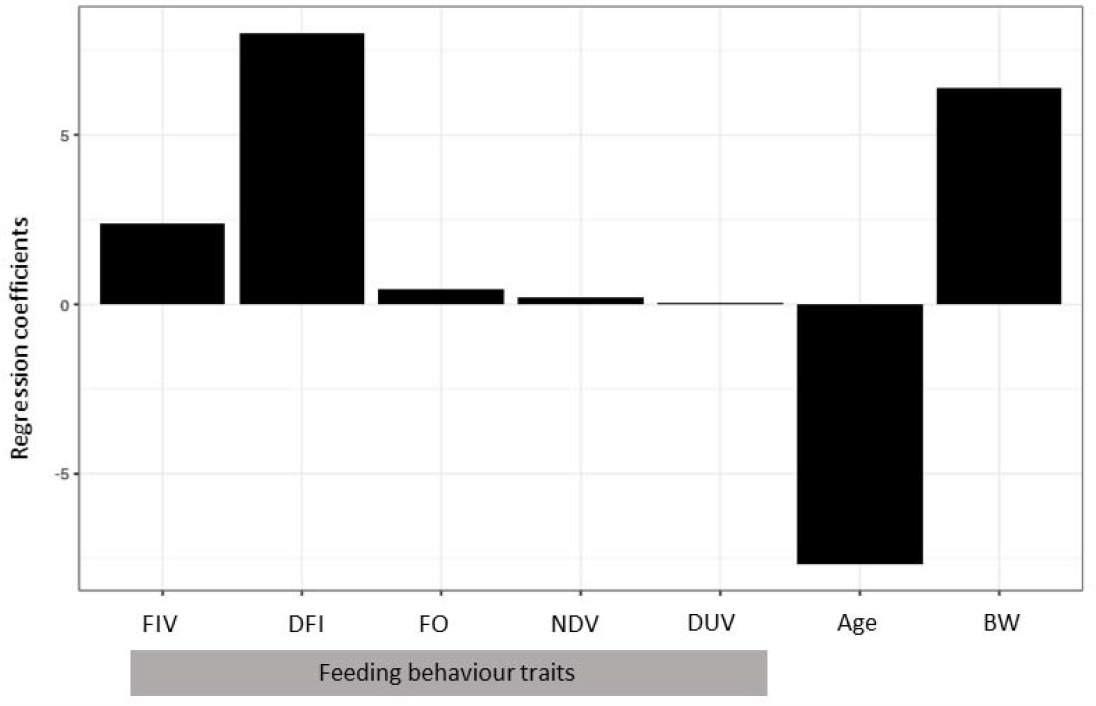
regression coefficient for daily feed intake, age at slaughter and body weight at slaughter and log-transformed regression coefficients for feed intake per visit, feeder occupation, number of daily visits and duration of visit from partial least square regression using feeding behaviour traits to predict lipid gain, and including the effects of age at slaughter and body weight at slaughter. FIV: feed intake per visit, DFI: daily feed intake, FO: feeder occupation, NDV: number of daily visits, DUV: duration of visit, Age: age at slaughter, BW: body weight at slaughter

The permutation strategy, as described in the *pls* package, was used to select the optimal number of components for the models. This strategy selects LV with minimum RMSE of prediction and fewer components while still maintaining the performance of the prediction model. The models with 5, 2, and 3 LV were chosen as the optimal model for predicting PE, EnE, and LipG, respectively, and were thereafter used in the PLS model to create a prediction equation. The chosen models explained 34.64%, 55.42% and 82.68% of the total variation observed in PE, EnE, and LipG, respectively (Table 3). The prediction accuracy (R^2^v) was higher for EnE and LipG than for PE (Table 4), which suggests that in pigs, EnE and LipG can be more accurately predicted from feeding behaviour traits compared to PE. The R^2^ validation (R^2^v) for the efficiency traits (PE, EnE and LipG) were slightly lower compared to the calibration (R^2^c) (Table 4). The CCC showed correlations significantly different from zero between predicted and observed values for PE, EnE, and LipG on the validation set (Table 4).

**Table 3.**
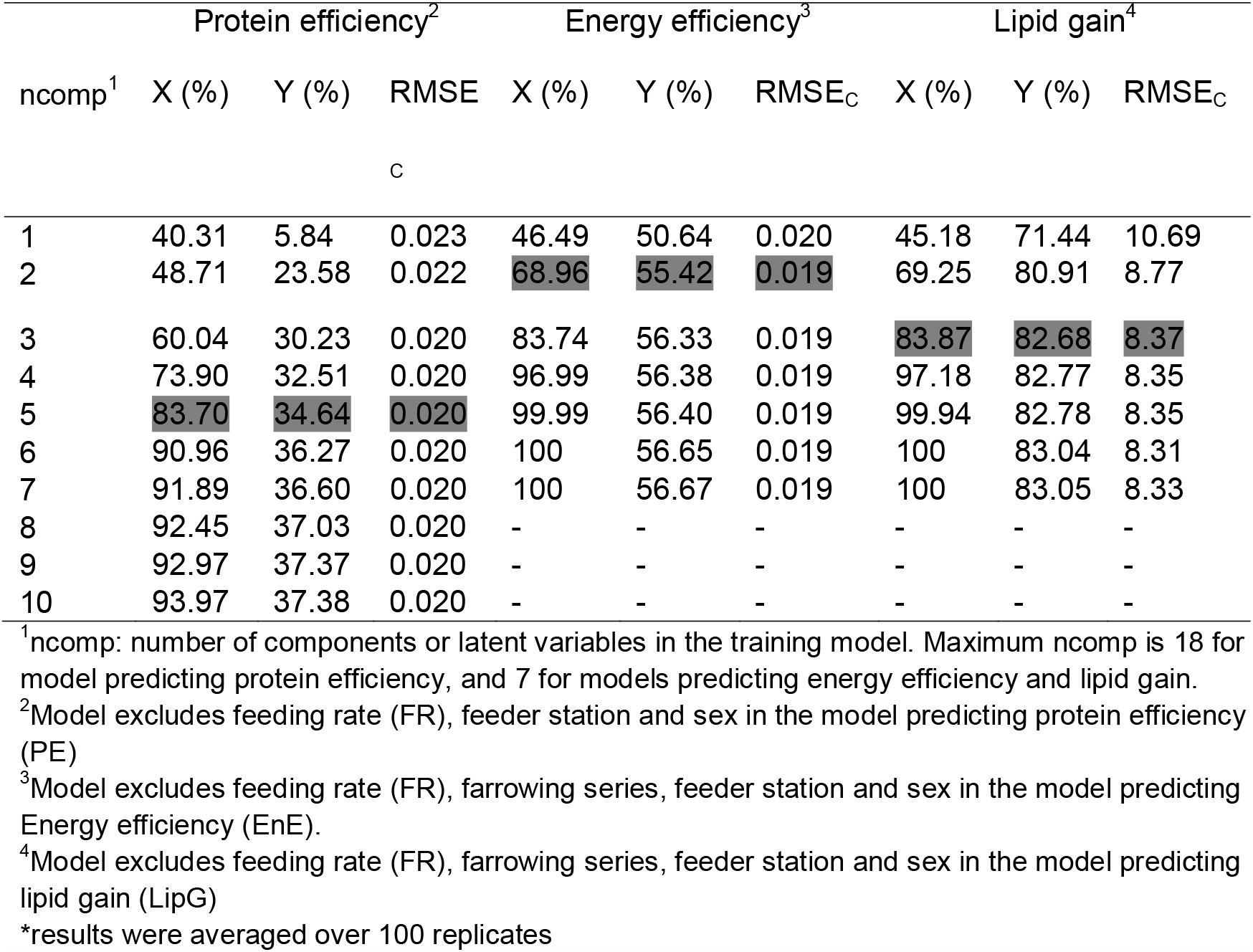
^*^Percentage of total variation explained in the independent (X; feeding behaviour traits, farrowing series, age and body weight at slaughter) and dependent variables (Y; protein efficiency, energy efficiency, and lipid gain), and the root mean square error (RMSE_C_) using partial least square regression on a calibration data set. The selected models with the optimal number of components are highlighted in grey.

**Table4.** ^*^Number of components, RMSE and R^2^ of the calibration and validation models in the partial least square regression analysis to predict protein efficiency (PE), energy efficiency (EnE) and daily lipid gain (LipG) using feeding behaviour traits. Prediction accuracy in terms of the mean bias and the concordance correlation coefficient are presented.

|  | Item | PE | EnE | LipG |
| --- | --- | --- | --- | --- |
| ncomp |  | 5 | 2 | 3 |
| Calibration | RMSE<sub>c</sub> | 0.020 | 0.019 | 8.37 |
| | $R^2_c$ (%) | 34.64 | 55.42 | 82.68 |
| Validation | RMSE<sub>v</sub> | 0.020 | 0.019 | 8.35 |
| | $R^2_v$ (%) | 32.86 | 53.76 | 82.41 |
| Prediction | Mean bias | 0.000011 | 0.000011 | 0.04 |
| accuracy | CCC | 0.50 [0.37, 0.60] | 0.70 [0.60, 0.77] | 0.90 [0.86, 0.93] |

$R^2_c$ = coefficient of determination for calibration; $R^2_v$ = coefficient of determination for validation; RMSE<sub>c</sub> = root mean square error of calibration; RMSE<sub>v</sub> = root mean square error of validation; CCC = concordance correlation coefficient
\*results were averaged over 100 replicates

## Discussion

Partial least square regression has been used in cattle to predict residual feed intake (RFI) from feeding behaviour traits with the aim of identifying and selecting efficient animals (Parsons 2018; Fischer et al., 2018). However, since energy is the main factor driving feed intake in pigs (Li and Patience, 2017), selection for improved RFI and feed conversion ratio (FCR) is a selection for energy efficiency rather than other specific nutrient efficiency, which may lead to an oversupply of nutrients (Kasper et al., 2020a; Millet et al., 2018). Moreover, considering the environmental impact associated with nitrogen excretion, selection for improved FCR and RFI is less efficient compared to selecting on the nutrient trait itself (de Verdal et al., 2011). Therefore, in this study, we investigated the potential of using feeding behaviour traits in predicting more specific efficiency traits (PE, EnE, and LipG) of Swiss Large White pigs when fed a protein-reduced diet. It is important to point out that the reduction in dietary protein content during the grower and finisher phase may have caused pigs to respond differently compared to when fed on usual protein recommendation. For instance, up to 19% reduction in the essential amino acid content of the diet was found to increase the feed intake of growing pigs but with no effects on the feed efficiency and the estimated protein retention compared to the pigs fed a standard diet (Schiavon et al., 2018).

### Protein efficiency

The result of this study showed that 34.64% of the variability in PE was explained by five feeding behaviour traits in addition to farrowing series, age and weight at slaughter, with negative relationships between PE and FIM, DFI and NDM. This suggests that protein-efficient pigs eat less per meal and visit the feeder less often. Ultimately, this explains the lower average daily feed intake observed in protein-efficient pigs. A strong positive association was observed between FR and protein retention in the study of Carcò et al. (2018), i.e., protein-efficient pigs eat at a faster rate, and FR was the most highly associated variable with protein and lipid gain. In contrast, the present data showed that FR was not important in the model (VIP < 1), and its inclusion did not improve the model in terms of R^2^c. For example, an exclusion of FR from the model in our study yielded an R^2^c of 32.51% and RMSE of 0.020 at component number of 4, while an inclusion of FR had R^2^c of 31.88% and RMSE of 0.020 at the same number of components. The reason for the difference in the study of Carcò et al. (2018) and our study may be due to differences in the weight and the period of protein restriction of the pigs. While pigs were fed a low-protein diet from an average weight of 22 kg to 106 kg BW in our study, Carcò et al. (2018) used pigs fed protein-reduced diet from 86 kg to 145 kg BW, a growth period where lipid deposition is much more important than protein retention. Another possible explanation to these discrepancies may be due to differences in the traits measured. Carcò et al. (2018) measured protein retention (g/day) while our study measured protein efficiency, which is the ratio between protein retention and protein consumed. Since PE is dependent on both, protein retained and consumed, an increase in protein retention does not necessarily mean an increase in PE, especially if the amount of protein consumed also increases when pigs are fed *ad libitum*. For example, a pig may consume X kg of dietary protein to make Y kg of body protein, while another pig may consume less dietary protein to make the same Y kg of body protein. In both cases, protein retention is the same, but protein efficiency is different. Similarly, Carcò et al. (2018) reported a positive partial correlation (r = 0.23) between DFI and protein retention, corrected for the effects of dietary treatments, while the slope from our study showed a significantly negative association between DFI and PE. This also may be due to differences in traits, as an increase in feed intake may truly increase protein retention but does not equate to a higher protein efficiency. To improve the prediction accuracy of PE, other predictors such as average daily gain, backfat thickness measured with ultrasound and carcass traits, could be included in future studies. For instance, the inclusion of initial backfat depth and gain in backfat depth in a PLS model to predict RFI by nine feeding behaviour traits in steers improved the percentage of variation explained by 3.9 % (Parsons et al., 2018). In the present study, while the use of only farrowing series, age at slaughter and body weight at slaughter in predicting PE had R^2^ validation of 11%, the addition of these variables to feeding behaviour traits increased R^2^ validation of PE from 4% to 32.86% (Supplementary Table S2).

### Energy efficiency and lipid gain

To our knowledge, there are no studies on the prediction of energy efficiency from feeding behaviour traits recorded by single-spaced automatic feeders. However, since selection for improved FCR and RFI is related to energy efficiency, we discuss studies on these classical efficiency traits, which were conducted in cattle. In our study, 55.42% of the variation in EnE was accounted for by five feeding behaviour traits in addition to age and weight at slaughter with positive relationships between EnE and FIM, DFI, FO and DUM. This indicates that energy efficient pigs visited the feeder more often and spent more time at the feeder, which ultimately resulted in a greater average daily feed intake. Feeding behaviour explained 21.3% of the variation in RFI in dairy cows (Fischer et al., 2018) and a negative relationship of RFI with duration of feeding and frequency of feeding per day in beef cattle (Durunna et al., 2011; Nkrumah et al., 2006). Similar to EnE, five feeding behaviour traits in addition to age and weight at slaughter explained 84.32% of the total variation observed in LipG. Although the use of body weight and age at slaughter to predict EnE and LipG gave R^2^ validation of 36% and 68%, respectively, the addition of these traits to feeding behaviour traits increased R^2^ validation of EnE and LipG from 41% and 65% to 53.76% and 82.41%, respectively (Supplementary Table S3 – S4). Positive relationships were observed between LipG and the feeding behaviour traits. This indicates that pigs that ate more, spent more time at the feeder, and visited the feeder more often gained more fat mass per day. However, these associations observed between LipG and feeding behaviour traits in our study differ from those reported by Carcò et al. (2018), where a negative relationship was found between daily lipid gain and feed intake and time spent eating. These differences may be due to factors such as breed, weight of pigs and the diet.

Overall, this study suggests that feeding behaviour traits, as well as age and body weight, can be used to predict PE, EnE, and LipG in Swiss Large White pigs under dietary protein restriction and *ad libitum* access to feed, but EnE and LipG can be predicted more accurately than PE. However, since these efficiency traits are complex, influenced by both genetic and environmental factors, feeding behaviour traits cannot be a sole indicator for these traits (Halachmi et al., 2016), especially for protein efficiency. Therefore, additional traits, such as carcass traits and average daily gain, may be included in the model to improve its accuracy and to capture more variation in the efficiency traits, especially for PE.

## Ethics approval

The experimental procedure was approved by the Office for Food Safety and Veterinary Affairs (2018_30_FR) and all procedures were conducted in accordance with the Ordinance on Animal Protection and the Ordinance on Animal Experimentation.

## Data and model availability statement

The data that support the findings of this study and the code used for models and the statistical analysis are publicly available in Zenodo (https://zenodo.org/record/5126661).

## Author contributions

Conceptualization: E.E., G.B. and C.K. Data Curation: E.E. Formal Analysis: E.E. Funding Acquisition: G.B. and C.K. Methodology: E.E. and C.K. Project Administration: C.K. Software: E.E. Supervision: G.B. and C.K. Visualization: E.E. Writing - Original Draft Preparation: E.E. Writing - Review & Editing: E.E., G.B. and C.K.

## Acknowledgements

We are grateful to Guy Maïkoff and his team for the maintenance and slaughter of the pigs and assistance with DXA scans.

