## Supplementary material for "Evaluation of feeding behaviour traits to predict efficiency traits in pigs using partial least square regression": Code: feedingbehaviour_PLSR.Rmd

````
---
title: "newfeedingbehaviour_PLSR"
author: "Ewaoluwagbemiga Esther"
date: "28/06/2021"
output: html_document
---

```{r setup, include=FALSE}
library(dplyr)
library(caret)
library(pls)
install.packages("termstrc", repos="http://R-Forge.R-project.org")
library('grid')
library(pls)
library(tidyverse)
library(egg)
library(DescTools)
library(SimDesign)
```


## import data

```{r cars}
behaviour_I <- read.table("feedingbehaviour.txt", header = TRUE, sep = "\t") 


boxplot(behaviour_I$prot_efficiency,
        ylab = "protein efficiency"
) # check for outliers using boxplot


behaviour1 <- behaviour_I[-c(192,547), ] #two pigs with high protein efficiency considered as outliers are removed

behaviour1$series <- as.factor(behaviour1$series)
behaviour1$station2 <- as.factor(behaviour1$station2)
behaviour1$sex <- as.factor(behaviour1$sex)

#log transformation
behaviour1$NDVvisits_day <- log10(behaviour1$NDVvisits_day) #logarithmic transformation of NDV
behaviour1$DUVmin_visit <- log10(behaviour1$DUVmin_visit) #logarithmic transformation of DUV
behaviour1$FOmin_day <- log10(behaviour1$FOmin_day) #logarithmic transformation of FO
behaviour1$FRg_min_day <- log10(behaviour1$FRg_min_day) #logarithmic transformation of FR
behaviour1$FIVg_visit <- log10(behaviour1$FIVg_visit) #logarithmic transformation of FIV

#Scale and center variables
behaviour1$DFIkg_day <- as.numeric(scale(behaviour1$DFIkg_day, center = TRUE))
behaviour1$NDVvisits_day <- as.numeric(scale(behaviour1$NDVvisits_day, center = TRUE))
behaviour1$FOmin_day <- as.numeric(scale(behaviour1$FOmin_day, center = TRUE))
behaviour1$FRg_min_day <- as.numeric(scale(behaviour1$FRg_min_day, center = TRUE))
behaviour1$FIVg_visit <- as.numeric(scale(behaviour1$FIVg_visit, center = TRUE))
behaviour1$DUVmin_visit <- as.numeric(scale(behaviour1$DUVmin_visit, center = TRUE))
behaviour1$age1 <- as.numeric(scale(behaviour1$age1, center = TRUE))
behaviour1$BWs <- as.numeric(scale(behaviour1$BWs, center = TRUE))


#relevel series
behaviour1 <- within(behaviour1, series <- relevel(series, ref = 12))
```

## subset data into training (calibration dataset) and test (validation dataset)

```{r }
indices <- caret::createDataPartition(behaviour1$prot_efficiency, p = 0.8, list = TRUE, times = 100)

resample_data <- tibble(
  training_sets = map(indices, ~ behaviour1[.x, ]),
  testing_sets = map(indices, ~ behaviour1[-.x, ])
)

```

## analysis for protein efficiency

## variable selection using the entire data set
```{r }
fullmodel_PE <- plsr(prot_efficiency ~ FRg_min_day + FIVg_visit + DFIkg_day + FOmin_day + NDVvisits_day + DUVmin_visit + age1 + series + station2 + sex + BWs, data = behaviour1, validation = "LOO", jackknife = TRUE)

summary(fullmodel_PE)

ncomp.permute <- selectNcomp(fullmodel_PE, method = "randomization", plot = TRUE,
                              ylim = c(0.019, 0.024)) #check for optimal number of components

#select variables with VIP > 1 at the optimal number of components.
VIP <- function(object) {
  if (object$method != "kernelpls")
    stop("Only implemented for orthogonal scores algorithm.  Refit with 'method = \"kernelpls\"'")
  if (nrow(object$Yloadings) > 1)
    stop("Only implemented for single-response models")
  
  SS <- c(object$Yloadings)^2 * colSums(object$scores^2)
  Wnorm2 <- colSums(object$loading.weights^2)
  SSW <- sweep(object$loading.weights^2, 2, SS / Wnorm2, "*")
  sqrt(nrow(SSW) * apply(SSW, 1, cumsum) / cumsum(SS))
}


vip_PE <- as.data.frame(VIP(fullmodel_PE))

vip_PE_comp7 <- vip_PE[5, ]

```


#analysis with variables of VIP > 1 in a final model using training dataset
```{r }
x_PE <- NULL
finalmodel_PE <- NULL

for (i in 1:nrow(resample_data)) {
finalmodel_PE[[i]] <- plsr(prot_efficiency ~ FIVg_visit + DFIkg_day + FOmin_day + NDVvisits_day + DUVmin_visit + age1 + series + BWs, data = resample_data[[1]][[i]], validation = "LOO", jackknife = TRUE)
x_PE[[i]] <-(capture.output(summary(finalmodel_PE[[i]])))
}

write.xlsx(x_PE, file = "PLS_PE.xlsx") 

for (i in 1:nrow(resample_data)) {plot(RMSEP(finalmodel_PE[[i]]), legendpos = "topright")} #plot RMSE 

 
for (i in 1:nrow(resample_data)) {ncomp.permute[[i]] <- selectNcomp(finalmodel_PE[[i]], method = "randomization", plot = TRUE, ylim = c(0.019, 0.024))}

write.xlsx(ncomp.permute, file = "ncomp_PE.xlsx")

for (i in 1:nrow(resample_data)) {plot(finalmodel_PE[[i]], ncomp = 5, asp = 1, line = TRUE)} #plot predicted versus measured values at optimal number of component.

jack.test(finalmodel_PE[[99]], ncomp=5) #jackknife test to obtain coefficients
```

#prediction and accuracy
```{r}
prediction_PE <- NULL

for (i in 1:nrow(resample_data)) {prediction_PE[[i]] <- predict(finalmodel_PE[[i]], ncomp = 5, newdata = resample_data[[2]][[i]]) }#predict protein efficiency from test data set

RMSE <- NULL
Rsquare <- NULL
for (i in 1:nrow(resample_data)) {
data.frame(Rsquare[[i]] <- caret::R2(prediction_PE[[i]], resample_data[[2]][[i]]$prot_efficiency),
  RMSE[[i]] <- caret::RMSE(prediction_PE[[i]], resample_data[[2]][[i]]$prot_efficiency) 
)
}#obtain RMSE and r-squared of prediction
write.xlsx(RMSE, file = "RMSE_PE.xlsx") #Save rmse values
write.xlsx(Rsquare, file = "rsquare_PE.xlsx") #save r-squared values

predict <- NULL

for (i in 1:nrow(resample_data)) {predict[[i]] <- as.data.frame(prediction_PE[[i]], row.names = NULL, optional = FALSE, 
              cut.names = FALSE, col.names = names(prediction), fix.empty.names = TRUE,
              stringsAsFactors = default.stringsAsFactors())}

predicted_PE <- NULL
for (i in 1:nrow(resample_data)) {predicted_PE[[i]] <- cbind(as.data.frame(resample_data[[2]][[i]]$ID), (predict[[i]]))}

for (i in 1:nrow(resample_data)) {names(predicted_PE[[i]]) <- c("ID", "prot_efficiency")}

#cor.test(predicted_PE$prot_efficiency, resample_data[[2]][[1]]$prot_efficiency, method = "spearman")

ccc <- NULL
for (i in 1:nrow(resample_data)) {ccc[[i]] <- CCC(predicted_PE[[i]]$prot_efficiency, resample_data[[2]][[i]]$prot_efficiency, ci = "z-transform",
               conf.level = 0.95)}

correlations <- NULL
for (i in 1:nrow(resample_data)) {correlations[[i]] <- ccc[[i]]$rho.c}

bias <- NULL
for (i in 1:nrow(resample_data)) {bias[[i]] <- bias(predicted_PE[[i]]$prot_efficiency, resample_data[[2]][[i]]$prot_efficiency)}

#rmse(predicted_PE$prot_efficiency, resample_data[[2]][[1]]$prot_efficiency)
write.xlsx(correlations, file = "correlation_PE.xlsx")
write.xlsx(bias, file = "bias_PE.xlsx")
```

```{r}

```

## subset data into training (calibration dataset) and test (validation dataset)

```{r }
indices <- caret::createDataPartition(behaviour1$energyefficiency, p = 0.8, list = TRUE, times = 100)

resample_data <- tibble(
  training_sets = map(indices, ~ behaviour1[.x, ]),
  testing_sets = map(indices, ~ behaviour1[-.x, ])
)

```

#analysis for energy efficiency
```{r}
fullmodel_EnE <- plsr(energyefficiency ~ FRg_min_day + FIVg_visit + DFIkg_day + FOmin_day + NDVvisits_day + DUVmin_visit + age1 + series + station2 + sex + BWs, data = behaviour1, validation = "LOO", jackknife = TRUE)

summary(fullmodel_EnE)

ncomp.permute <- selectNcomp(fullmodel_EnE, method = "randomization", plot = TRUE,
                              ylim = c(0.017, 0.029)) #check for optimal number of components

#select variables with VIP > 1 at the optimal number of components.
VIP <- function(object) {
  if (object$method != "kernelpls")
    stop("Only implemented for orthogonal scores algorithm.  Refit with 'method = \"kernelpls\"'")
  if (nrow(object$Yloadings) > 1)
    stop("Only implemented for single-response models")
  
  SS <- c(object$Yloadings)^2 * colSums(object$scores^2)
  Wnorm2 <- colSums(object$loading.weights^2)
  SSW <- sweep(object$loading.weights^2, 2, SS / Wnorm2, "*")
  sqrt(nrow(SSW) * apply(SSW, 1, cumsum) / cumsum(SS))
}


vip_EnE <- as.data.frame(VIP(fullmodel_EnE))

vip_EnE_comp7 <- vip_EnE[7, ]

```

#analysis with variables of VIP > 1 in a final model using training dataset
```{r }
x_EnE <- NULL
finalmodel_EnE <- NULL

for (i in 1:nrow(resample_data)) {
finalmodel_EnE[[i]] <- plsr(energyefficiency ~ FIVg_visit + DFIkg_day + FOmin_day + NDVvisits_day + DUVmin_visit + age1 + BWs, data = resample_data[[1]][[i]], validation = "LOO", jackknife = TRUE)
x_EnE[[i]] <-(capture.output(summary(finalmodel_EnE[[i]])))
}

write.xlsx(x_EnE, file = "PLS_EnE.xlsx") 

for (i in 1:nrow(resample_data)) {plot(RMSEP(finalmodel_EnE[[i]]), legendpos = "topright")} #plot RMSE 

 
for (i in 1:nrow(resample_data)) {ncomp.permute[[i]] <- selectNcomp(finalmodel_EnE[[i]], method = "randomization", plot = TRUE, ylim = c(0.018, 0.030))}

write.xlsx(ncomp.permute, file = "ncomp_EnE.xlsx")

for (i in 1:nrow(resample_data)) {plot(finalmodel_EnE[[i]], ncomp = 2, asp = 1, line = TRUE)} #plot predicted versus measured values at optimal number of component.

jack.test(finalmodel_PE[[99]], ncomp=2) #jackknife test to obtain coefficients

```

#prediction and accuracy
```{r}
prediction_EnE <- NULL

for (i in 1:nrow(resample_data)) {prediction_EnE[[i]] <- predict(finalmodel_EnE[[i]], ncomp = 2, newdata = resample_data[[2]][[i]]) }#predict protein efficiency from test data set

RMSE <- NULL
Rsquare <- NULL
for (i in 1:nrow(resample_data)) {
data.frame(Rsquare[[i]] <- caret::R2(prediction_EnE[[i]], resample_data[[2]][[i]]$energyefficiency),
  RMSE[[i]] <- caret::RMSE(prediction_EnE[[i]], resample_data[[2]][[i]]$energyefficiency) 
)
}#obtain RMSE and r-squared of prediction

write.xlsx(RMSE, file = "RMSE_EnE.xlsx")
write.xlsx(Rsquare, file = "rsquare_EnE.xlsx")

predict <- NULL

for (i in 1:nrow(resample_data)) {predict[[i]] <- as.data.frame(prediction_EnE[[i]], row.names = NULL, optional = FALSE, 
              cut.names = FALSE, col.names = names(prediction), fix.empty.names = TRUE,
              stringsAsFactors = default.stringsAsFactors())}

predicted_EnE <- NULL
for (i in 1:nrow(resample_data)) {predicted_EnE[[i]] <- cbind(as.data.frame(resample_data[[2]][[i]]$ID), (predict[[i]]))}

for (i in 1:nrow(resample_data)) {names(predicted_EnE[[i]]) <- c("ID", "energyefficiency")}

#cor.test(predicted_PE$prot_efficiency, resample_data[[2]][[1]]$prot_efficiency, method = "spearman")

ccc <- NULL
for (i in 1:nrow(resample_data)) {ccc[[i]] <- CCC(predicted_EnE[[i]]$energyefficiency, resample_data[[2]][[i]]$energyefficiency, ci = "z-transform",
               conf.level = 0.95)}

correlations <- NULL
for (i in 1:nrow(resample_data)) {correlations[[i]] <- ccc[[i]]$rho.c}

bias <- NULL
for (i in 1:nrow(resample_data)) {bias[[i]] <- bias(predicted_EnE[[i]]$energyefficiency, resample_data[[2]][[i]]$energyefficiency)}

#rmse(predicted_PE$prot_efficiency, resample_data[[2]][[1]]$prot_efficiency)
write.xlsx(correlations, file = "correlation_EnE.xlsx")
write.xlsx(bias, file = "bias_EnE.xlsx")

```

```{r}

```
## subset data into training (calibration dataset) and test (validation dataset)

```{r }
indices <- caret::createDataPartition(behaviour1$fatgain, p = 0.8, list = TRUE, times = 100)

resample_data <- tibble(
  training_sets = map(indices, ~ behaviour1[.x, ]),
  testing_sets = map(indices, ~ behaviour1[-.x, ])
)

```


# analysis for lipid gain
```{r}
fullmodel_LipG <- plsr(fatgain ~ FRg_min_day + FIVg_visit + DFIkg_day + FOmin_day + NDVvisits_day + DUVmin_visit + age1 + series + station2 + sex + BWs, data = behaviour1, validation = "LOO", jackknife = TRUE)

summary(fullmodel_LipG)

ncomp.permute <- selectNcomp(fullmodel_LipG, method = "randomization", plot = TRUE,
                              ylim = c(7, 20)) #check for optimal number of components

#select variables with VIP > 1 at the optimal number of components.
VIP <- function(object) {
  if (object$method != "kernelpls")
    stop("Only implemented for orthogonal scores algorithm.  Refit with 'method = \"kernelpls\"'")
  if (nrow(object$Yloadings) > 1)
    stop("Only implemented for single-response models")
  
  SS <- c(object$Yloadings)^2 * colSums(object$scores^2)
  Wnorm2 <- colSums(object$loading.weights^2)
  SSW <- sweep(object$loading.weights^2, 2, SS / Wnorm2, "*")
  sqrt(nrow(SSW) * apply(SSW, 1, cumsum) / cumsum(SS))
}


vip_LipG <- as.data.frame(VIP(fullmodel_LipG))

vip_LipG_comp8 <- vip_LipG[8, ]

```

#analysis with variables of VIP > 1 in a final model using training dataset
```{r }
x_LipG <- NULL
finalmodel_LipG <- NULL

for (i in 1:nrow(resample_data)) {
finalmodel_LipG[[i]] <- plsr(fatgain ~ FIVg_visit + DFIkg_day + FOmin_day + NDVvisits_day + DUVmin_visit + age1 + BWs, data = resample_data[[1]][[i]], validation = "LOO", jackknife = TRUE)
x_LipG[[i]] <-(capture.output(summary(finalmodel_LipG[[i]])))
}

write.xlsx(x_LipG, file = "PLS_LipG.xlsx") 

for (i in 1:nrow(resample_data)) {plot(RMSEP(finalmodel_LipG[[i]]), legendpos = "topright")} #plot RMSE 

 
for (i in 1:nrow(resample_data)) {ncomp.permute[[i]] <- selectNcomp(finalmodel_LipG[[i]], method = "randomization", plot = TRUE, ylim = c(7, 20))}

write.xlsx(ncomp.permute, file = "ncomp_LipG.xlsx")

for (i in 1:nrow(resample_data)) {plot(finalmodel_LipG[[i]], ncomp = 3, asp = 1, line = TRUE)} #plot predicted versus measured values at optimal number of component.

jack.test(finalmodel_PE[[99]], ncomp=3) #jackknife test to obtain coefficients

```

#prediction and accuracy
```{r}
prediction_LipG <- NULL

for (i in 1:nrow(resample_data)) {prediction_LipG[[i]] <- predict(finalmodel_LipG[[i]], ncomp = 3, newdata = resample_data[[2]][[i]]) }#predict protein efficiency from test data set

RMSE <- NULL
Rsquare <- NULL
for (i in 1:nrow(resample_data)) {
data.frame(Rsquare[[i]] <- caret::R2(prediction_LipG[[i]], resample_data[[2]][[i]]$fatgain),
  RMSE[[i]] <- caret::RMSE(prediction_LipG[[i]], resample_data[[2]][[i]]$fatgain) 
)
}#obtain RMSE and r-squared of prediction

write.xlsx(RMSE, file = "RMSE_LipG.xlsx")
write.xlsx(Rsquare, file = "rsquare_LipG.xlsx")

predict <- NULL

for (i in 1:nrow(resample_data)) {predict[[i]] <- as.data.frame(prediction_LipG[[i]], row.names = NULL, optional = FALSE, 
              cut.names = FALSE, col.names = names(prediction), fix.empty.names = TRUE,
              stringsAsFactors = default.stringsAsFactors())}

predicted_LipG <- NULL
for (i in 1:nrow(resample_data)) {predicted_LipG[[i]] <- cbind(as.data.frame(resample_data[[2]][[i]]$ID), (predict[[i]]))}

for (i in 1:nrow(resample_data)) {names(predicted_LipG[[i]]) <- c("ID", "fatgain")}

#cor.test(predicted_PE$prot_efficiency, resample_data[[2]][[1]]$prot_efficiency, method = "spearman")

ccc <- NULL
for (i in 1:nrow(resample_data)) {ccc[[i]] <- CCC(predicted_LipG[[i]]$fatgain, resample_data[[2]][[i]]$fatgain, ci = "z-transform",
               conf.level = 0.95)}

correlations <- NULL
for (i in 1:nrow(resample_data)) {correlations[[i]] <- ccc[[i]]$rho.c}

bias <- NULL
for (i in 1:nrow(resample_data)) {bias[[i]] <- bias(predicted_LipG[[i]]$fatgain, resample_data[[2]][[i]]$fatgain)}

#rmse(predicted_PE$prot_efficiency, resample_data[[2]][[1]]$prot_efficiency)
write.xlsx(correlations, file = "correlation_LipG.xlsx")
write.xlsx(bias, file = "bias_LipG.xlsx")


```

# coefficient plot for protein efficiency
```{r}
df_coef <- NULL
for (i in 1:nrow(resample_data)) {df_coef[[i]] <- as.data.frame(coef(finalmodel_PE[[i]], ncomp = 5))}

write.xlsx(df_coef, file = "df_coef.xlsx")

df_coef_PE <- read.table("~/mnt/Data-Work-PO/23_Livestock-PO/233.1_pig_production/PigEfficiency/Heritability_Protein_Efficiency/Esther_Ewaoluwagbemiga/feedingbehaviour_animal/revised/Analysis/df_coef_PE.txt", header = TRUE, sep = "\t")

df_coef_PE <- column_to_rownames(df_coef_PE, var = "var")

df_coef_PE <- df_coef_PE %>% 
  dplyr::mutate(variables = rownames(.)) %>% 
  dplyr::mutate(variables = factor(variables, 
                                   levels = c(
                                     "FIVg_visit", "DFIkg_day", "FOmin_day", "NDVvisits_day", "DUVmin_visit", "age1", "series1", "series2", "series3", "series4", "series5", "series6", "series7", "series8", "series9", "series10", "series11", "BWs")
  ))

colnames(df_coef_PE)[1] <- "regression_coefficients"

df_coef_PE

p2_coef <-
  ggplot(df_coef_PE, aes(x = variables, y = regression_coefficients, group = 1))+  geom_bar(stat = "identity",  fill = "black")+
  theme(axis.text.x = element_text(angle=65,
                                   hjust=1,
                                   size = 8),
        axis.title.y = element_text(size = 2))+
  theme_bw()+
  theme(axis.text.x = element_blank())+
  labs(x="")


p_fin <- grid.draw(egg::ggarrange(p2_coef , ncol=1))
```

#coefficient plot for energy efficiency
```{r}
df_coef <- NULL
for (i in 1:nrow(resample_data)) {df_coef[[i]] <- as.data.frame(coef(finalmodel_EnE[[i]], ncomp = 2))}

write.xlsx(df_coef, file = "df_coef.xlsx")

df_coef_EnE <- read.table("~/mnt/Data-Work-PO/23_Livestock-PO/233.1_pig_production/PigEfficiency/Heritability_Protein_Efficiency/Esther_Ewaoluwagbemiga/feedingbehaviour_animal/revised/Analysis/df_coef_EnE.txt", header = TRUE, sep = "\t")

df_coef_EnE <- column_to_rownames(df_coef_EnE, var = "var")

df_coef_EnE <- df_coef_EnE %>% 
  dplyr::mutate(variables = rownames(.)) %>% 
  dplyr::mutate(variables = factor(variables, 
                                   levels = c(
                                     "FIVg_visit", "DFIkg_day", "FOmin_day", "NDVvisits_day", "DUVmin_visit", "age1", "BWs")
  ))

colnames(df_coef_EnE)[1] <- "regression_coefficients"

df_coef_EnE

p2_coef <-
  ggplot(df_coef_EnE, aes(x = variables, y = regression_coefficients, group = 1))+  geom_bar(stat = "identity",  fill = "black")+
  theme(axis.text.x = element_text(angle=65,
                                   hjust=1,
                                   size = 8),
        axis.title.y = element_text(size = 2))+
  theme_bw()+
  theme(axis.text.x = element_blank())+
  labs(x="")


p_fin <- grid.draw(egg::ggarrange(p2_coef , ncol=1))
```

#coefficient plot for fatgain
```{r}
df_coef <- NULL
for (i in 1:nrow(resample_data)) {df_coef[[i]] <- as.data.frame(coef(finalmodel_LipG[[i]], ncomp = 3))}

write.xlsx(df_coef, file = "df_coef.xlsx")

df_coef_LipG <- read.table("~/mnt/Data-Work-PO/23_Livestock-PO/233.1_pig_production/PigEfficiency/Heritability_Protein_Efficiency/Esther_Ewaoluwagbemiga/feedingbehaviour_animal/revised/Analysis/df_coef_LipG.txt", header = TRUE, sep = "\t")

df_coef_LipG <- column_to_rownames(df_coef_LipG, var = "var")

df_coef_LipG <- df_coef_LipG %>% 
  dplyr::mutate(variables = rownames(.)) %>% 
  dplyr::mutate(variables = factor(variables, 
                                   levels = c(
                                     "FIVg_visit", "DFIkg_day", "FOmin_day", "NDVvisits_day", "DUVmin_visit", "age1", "BWs")
  ))

colnames(df_coef_LipG)[1] <- "regression_coefficients"

df_coef_LipG

p2_coef <-
  ggplot(df_coef_LipG, aes(x = variables, y = regression_coefficients, group = 1))+  geom_bar(stat = "identity",  fill = "black")+
  theme(axis.text.x = element_text(angle=65,
                                   hjust=1,
                                   size = 8),
        axis.title.y = element_text(size = 2))+
  theme_bw()+
  theme(axis.text.x = element_blank())+
  labs(x="")


p_fin <- grid.draw(egg::ggarrange(p2_coef , ncol=1))
```

#pearson correlation between efficiency and feeding behaviour traits
```{r}
cor.test(x=behaviour1$FIVg_visit, y=behaviour1$prot_efficiency, method = "pearson")
cor.test(x=behaviour1$DFIg_day, y=behaviour1$prot_efficiency, method = "pearson")
cor.test(x=behaviour1$FOmin_day, y=behaviour1$prot_efficiency, method = "pearson")
cor.test(x=behaviour1$NDVvisits_day, y=behaviour1$prot_efficiency, method = "pearson")
cor.test(x=behaviour1$DUVmin_visit, y=behaviour1$prot_efficiency, method = "pearson")
cor.test(x=behaviour1$FRg_min_day, y=behaviour1$prot_efficiency, method = "pearson")

cor.test(x=behaviour1$FIVg_visit, y=behaviour1$prot_efficiency, method = "pearson")
cor.test(x=behaviour1$DFIg_day, y=behaviour1$prot_efficiency, method = "pearson")
cor.test(x=behaviour1$FOmin_day, y=behaviour1$prot_efficiency, method = "pearson")
cor.test(x=behaviour1$NDVvisits_day, y=behaviour1$prot_efficiency, method = "pearson")
cor.test(x=behaviour1$DUVmin_visit, y=behaviour1$prot_efficiency, method = "pearson")
cor.test(x=behaviour1$FRg_min_day, y=behaviour1$prot_efficiency, method = "pearson")

cor.test(x=behaviour1$NDVvisits_day, y=behaviour1$FRg_min_day, method = "pearson")
cor.test(x=behaviour1$DUVmin_visit, y=behaviour1$FRg_min_day, method = "pearson")
cor.test(x=behaviour1$FIVg_visit, y=behaviour1$FRg_min_day, method = "pearson")
cor.test(x=behaviour1$DFIg_day, y=behaviour1$FRg_min_day, method = "pearson")
cor.test(x=behaviour1$FOmin_day, y=behaviour1$FRg_min_day, method = "pearson")
cor.test(x=behaviour1$DFIg_day, y=behaviour1$FIVg_visit, method = "pearson")
cor.test(x=behaviour1$FOmin_day, y=behaviour1$FIVg_visit, method = "pearson")
cor.test(x=behaviour1$DUVmin_visit, y=behaviour1$FIVg_visit, method = "pearson")
cor.test(x=behaviour1$NDVvisits_day, y=behaviour1$FIVg_visit, method = "pearson")
cor.test(x=behaviour1$FOmin_day, y=behaviour1$DFIg_day, method = "pearson")
cor.test(x=behaviour1$DUVmin_visit, y=behaviour1$DFIg_day, method = "pearson")
cor.test(x=behaviour1$NDVvisits_day, y=behaviour1$DFIg_day, method = "pearson")
cor.test(x=behaviour1$DUVmin_visit, y=behaviour1$FOmin_day, method = "pearson")
cor.test(x=behaviour1$NDVvisits_day, y=behaviour1$FOmin_day, method = "pearson")
cor.test(x=behaviour1$NDVvisits_day, y=behaviour1$DUVmin_visit, method = "pearson")

```
```{r pressure, echo=FALSE}
plot(pressure)
```

Note that the `echo = FALSE` parameter was added to the code chunk to prevent printing of the R code that generated the plot.
````
