## Supplementary tables for "Evaluation of feeding behaviour traits to predict efficiency traits in pigs using partial least square regression"

Table S1: R-squared calibration and validation obtained from different statistical methods and the correlations of predictions between PLSR and Bayesian models (CorrP) for the efficiency traits

| Traits* | PLSR | | BRR | | BayesA | | BayesB | | CorrP | | |
| --- | --- | --- | --- | --- | --- | --- | --- | --- | --- | --- | --- |
|  | R^2^c | R^2^v | R^2^c | R^2^v | R^2^c | R^2^v | R^2^c | R^2^v | BRR | BayesA | BayesB |
| Protein efficiency | 0.35 | 0.33 | 0.4 | 0.35 | 0.37 | 0.33 | 0.36 | 0.32 | 0.97 | 0.96 | 0.98 |
| Energy efficiency | 0.55 | 0.54 | 0.56 | 0.56 | 0.58 | 0.48 | 0.58 | 0.48 | 0.99 | 0.99 | 0.99 |
| Lipid gain | 0.83 | 0.82 | 0.84 | 0.83 | 0.83 | 0.82 | 0.83 | 0.82 | 1.00 | 1.00 | 1.00 |

R^2^c = R-squared calibration; R^2^v = R-squared validation; PLSR = partial least square regression; BRR = Bayesian ridge regression. R-squared is the squared correlations between observed and predicted values for the validation dataset. CorrP is the correlations between PLSR and Bayesian models for the same individuals in the validation dataset.

*Results were averaged over 25 replicates using 6000 iterations and 1000 burn-ins.

|  |  | | Protein efficiency (PE) | | | | | | | | |
| --- | --- | --- | --- | --- | --- | --- | --- | --- | --- | --- | --- |
|  |  | | Model A^1^ | | | Model B^2^ | | | Model C | | |
|  |  | | X (%) | Y (%) | RMSE | X (%) | Y (%) | RMSE | X (%) | Y (%) | RMSE |
|  | ncomp | |  |  |  |  |  |  |  |  |  |
| Calibration | 1 | | 40.31 | 5.84 | 0.023 | 33.21 | 4.34 | 0.024 | 60.25 | 3.76 | 0.023 |
|  | 2 | | 48.71 | 23.58 | 0.022 | 40.69 | 15.52 | 0.023 | 83.83 | 5.14 | 0.023 |
|  | 3 | | 60.04 | 30.23 | 0.02 | 74.14 | 15.93 | 0.023 | 99.99 | 5.57 | 0.023 |
|  | 4 | | 73.9 | 32.51 | 0.02 | 76.66 | 16.7 | 0.023 | 100 | 8.71 | 0.023 |
|  | 5 | | 83.7 | 34.64 | 0.02 | 77.24 | 19.1 | 0.023 | 100 | 8.77 | 0.023 |
|  | 6 | | 90.96 | 36.27 | 0.02 | 80.19 | 19.23 | 0.003 |  |  |  |
|  | 7 | | 91.89 | 36.6 | 0.02 | 83.21 | 19.23 | 0.003 |  |  |  |
|  | 8 | | 92.45 | 37.03 | 0.02 | 85.85 | 19.23 | 0.003 |  |  |  |
|  | 9 | | 92.97 | 37.37 | 0.02 | 88.67 | 19.23 | 0.003 |  |  |  |
|  | 10 | | 93.97 | 37.38 | 0.02 | 91.29 | 19.23 | 0.003 |  |  |  |
| ***Validation** | |  | **-** | **32.86** | **0.02** | **-** | **11** | **0.022** | **-** | **4** | **0.024** |

Table S2: percentage of total variation explained in the predictors (X) and response variable (Y; protein efficiency) and root mean square error (RMSE) between models for calibration and validation data set using partial least square regression.

*The selected models with the optimal number of components used on the validation sets are highlighted in grey.

Model A includes duration per meal (DUM), feed intake per meal (FIM), number of daily meals (NDM), feeder occupation (FO), daily feed intake (DFI), farrowing series, age at slaughter and body weight as slaughter as the predictors.

Model B includes farrowing series, age at slaughter and body weight at slaughter as predictors.

Model C includes duration per meal (DUM), feed intake per meal (FIM), number of daily meals (NDM), feeder occupation (FO) and daily feed intake (DFI).

^1^Maximum number of components (ncomp) is 18; ^2^Maximum number of components (ncomp) is 13.

|  |  | | Energy efficiency (EnE) | | | | | | | | |
| --- | --- | --- | --- | --- | --- | --- | --- | --- | --- | --- | --- |
|  |  | | Model A | | | Model B | | | Model C | | |
|  |  | | X (%) | Y (%) | RMSE | X (%) | Y (%) | RMSE | X (%) | Y (%) | RMSE |
|  | ncomp | |  |  |  |  |  |  |  |  |  |
| Calibration | 1 | | 46.49 | 50.64 | 0.02 | 49.12 | 47.59 | 0.021 | 60.45 | 39.48 | 0.023 |
|  | 2 | | 68.96 | 55.42 | 0.019 | 100 | 47.75 | 0.021 | 87.12 | 46.53 | 0.021 |
|  | 3 | | 83.74 | 56.33 | 0.019 |  |  |  | 99.99 | 49.45 | 0.021 |
|  | 4 | | 96.99 | 56.38 | 0.019 |  |  |  | 100 | 51.42 | 0.020 |
|  | 5 | | 99.99 | 56.4 | 0.019 |  |  |  | 100 | 51.43 | 0.020 |
|  | 6 | | 100 | 56.65 | 0.019 |  |  |  |  |  |  |
|  | 7 | | 100 | 56.67 | 0.019 |  |  |  |  |  |  |
| ***Validation** | |  | **-** | **53.76** | **0.019** | **-** | **36** | **0.022** | **-** | **41** | **0.02** |

Table S3: percentage of total variation explained in the predictors (X) and response variable (Y; energy efficiency) and root mean square error (RMSE) between models for calibration and validation data set using partial least square regression.

*The selected models with the optimal number of components used on the validation sets are highlighted in grey.

Model A includes duration per meal (DUM), feed intake per meal (FIM), number of daily meals (NDM), feeder occupation (FO), daily feed intake (DFI), age at slaughter and body weight at slaughter as predictors.

Model B includes age at slaughter and body weight at slaughter as predictors.

Model C includes duration per meal (DUM), feed intake per meal (FIM), number of daily meals (NDM), feeder occupation (FO) and daily feed intake (DFI).

Table S4: percentage of total variation explained in the predictors (X) and response variable (Y; lipid gain) and root mean square error (RMSE) between models for calibration and validation data set using partial least square regression.

|  |  | | Lipid gain (LipG) | | | | | | | | |
| --- | --- | --- | --- | --- | --- | --- | --- | --- | --- | --- | --- |
|  |  | | Model A | | | Model B | | | Model C | | |
|  |  | | X (%) | Y (%) | RMSE | X (%) | Y (%) | RMSE | X (%) | Y (%) | RMSE |
|  | ncomp | |  |  |  |  |  |  |  |  |  |
| Calibration | 1 | | 45.18 | 71.44 | 10.69 | 48.91 | 72.86 | 10.44 | 56.9 | 56.06 | 12.95 |
|  | 2 | | 69.25 | 80.91 | 8.77 | 100 | 73.09 | 10.39 | 86.74 | 67.9 | 11.09 |
|  | 3 | | 83.87 | 82.68 | 8.37 |  |  |  | 99.99 | 72.99 | 10.17 |
|  | 4 | | 97.18 | 82.77 | 8.35 |  |  |  | 100 | 73.7 | 10.05 |
|  | 5 | | 99.94 | 82.78 | 8.35 |  |  |  | 100 | 73.71 | 10.08 |
|  | 6 | | 100 | 83.04 | 8.31 |  |  |  |  |  |  |
|  | 7 | | 100 | 83.05 | 8.33 |  |  |  |  |  |  |
| ***Validation** | |  | **-** | **82.41** | **8.35** | **-** | **68** | **11.29** | **-** | **65** | **12.71** |

*The selected models with the optimal number of components used on the validation sets are highlighted in grey.

Model A includes duration per meal (DUM), feed intake per meal (FIM), number of daily meals (NDM), feeder occupation (FO), daily feed intake (DFI), age at slaughter and body weight at slaughter as predictors.

Model B includes age at slaughter and body weight at slaughter as predictors.

Model C includes duration per meal (DUM), feed intake per meal (FIM), number of daily meals (NDM), feeder occupation (FO) and daily feed intake (DFI).
